# An atlas of *Brachypodium distachyon* lateral root development

**DOI:** 10.1101/2024.03.25.586236

**Authors:** Cristovao De Jesus Vieira Teixeira, Kevin Bellande, Alja van der Schuren, Devin O’Connor, Christian S. Hardtke, Joop EM Vermeer

**Affiliations:** Laboratory of Molecular and Cell Biology, Institute of Biology, University of Neuchâtel, Switzerland; Department of Plant Molecular Biology, University of Lausanne, Lausanne, Switzerland; IPSiM, University of Montpellier, CNRS, INRAE, Institut Agro, Montpellier, France; Sainsbury Lab, University of Cambridge, Cambridge, U.K

## Abstract

The root system of plants is a vital part for successful development and adaptation to different soil types and environments. Besides allowing exploration of the soil for water and nutrients, it also provides anchorage. A major determinant of the shape of a plant root system is the formation of lateral roots, allowing for expansion of the root system. *Arabidopsis thaliana*, with its simple root anatomy, has been extensively studied to reveal the genetic program underlying root branching. However, to get a more general understanding of lateral root development, comparative studies in species with a more complex root anatomy are required. *Brachypodium distachyon* is a wild, temperate grass species, that is related to important crops such as wheat. Its roots contain multiple cortex layers and an exodermis that functions as an additional root barrier, besides the endodermis. Here, by combining optimized clearing methods and histology, we describe an atlas of lateral root development in Brachypodium. We show that lateral roots initiate from enlarged phloem pole pericycle cells and that the overlying endodermis reactivates its cell cycle and eventually forms the root cap. In addition, auxin signaling reported by the DR5 reporter was not detected in the phloem pole pericycle cells or young primordia. In contrast, auxin signaling was activated in the overlying cell cortical layers, including the exodermis. Thus, Brachypodium is a valuable model to investigate how signaling pathways and cellular responses have been repurposed to facilitate lateral root organogenesis.

## Introduction

Root branching is vital for plant survival as it facilitates the uptake of water and nutrients (Orosa-Puente *et al*., 2018). Root system architecture (RSA) consists of structural features like root length, spread, number, and length of lateral roots (LRs), among others (Bao *et al*., 2014; Morris *et al*., 2017; Kumar *et al*., 2019). RSA exhibits great plasticity in response to environmental changes and it is a desirable trait to breed more resilient crops (Ye *et al*., 2017; Yu, Hochholdinger and Li, 2019; Schäfer *et al*., 2022). In both monocots and dicots, the growth angle and number of LRs are the central components of the overall RSA (Atkinson *et al*., 2014; Roychoudhry *et al*., 2017). However, the molecular and cell biological program underlying root branching are less described for major crops due to the difficulty of observing the root system throughout the plant’s life cycle (Hochholdinger and Zimmermann, 2008).

Due to the relatively simple organization of its root system, tissue transparency and extensive genetic tool box, *Arabidopsis thaliana* (Arabidopsis) has been the most characterized experimental system for dissecting the molecular mechanisms underlying LR development (Banda *et al*., 2019). In Arabidopsis, LRs initiate from Lateral Root Founder Cells (LRFCs), and a series of highly coordinated cell divisions leads to the development of a new LR primordium (LRP) (Casimiro *et al*., 2001; Ditengou *et al*., 2008; Stoeckle, Thellmann and Vermeer, 2018; Gala *et al*., 2021). In this case, LRFCs are patterned along the primary root axis with a regulated spacing, beginning from the basal root meristem (Lavenus *et al*., 2015; Chen *et al*., 2018; Kircher and Schopfer, 2018; Torres-Martínez *et al*., 2020). Early stage LRP are more likely initiated closer to the root tip. The first morphological event of LR initiation takes place in the differentiation zone where LRFC founders cells divide asymmetrically and anticlinal (Malamy and Benfey, 1997). In addition, auxin signaling in the neighboring endodermis plays a major role during LRP formation as blocking auxin responses in this tissue abolishes LR formation (Vermeer, et al., 2014). Subsequent periclinal and anticlinal divisions give rise to an organized dome shaped LRP (Malamy and Benfey, 1997).

In monocots, LR studies have mostly been conducted on rice and maize (Wang *et al*., 2002; Hochholdinger and Zimmermann, 2008; Jansen *et al*., 2012; Uga *et al*., 2013). Notably, LR initiation in monocots predominantly occurs in the phloem-associated pericycle (Jansen et al., 2013; Hardtke and Pacheco-Villalobos, 2015) and the underlying mechanisms governing the patterning of LR formation in these agriculturally important crops are not well described. In contrast to Arabidopsis, in monocots and many other plant species, during LR development both the pericycle and endodermis undergo cell divisions, thereby contributing to the formation of the new organ (Casero, Casimiro and Lloret, 1995; Rebouillat *et al*., 2009; Banda *et al*., 2019; Xiao *et al*., 2019). However, only few studies have investigated the auxin-mediated transcriptome changes underlying LRP formation in monocots (Stelpflug *et al*., 2016; Kortz, Hochholdinger and Yu, 2019). Moreover, it is still unknown which signal is regulating the cell divisions in the endodermis overlying the newly formed LR.

The endodermis is the innermost cortical cell layer surrounding the vasculature (Geldner, 2013). Casparian strips (CS) and suberin lamellae (SL) formed in this layer were shown to be crucial in regulating the uptake of nutrients, in the response to osmotic stress and protection against pathogens (Ranathunge *et al*., 2008; Barberon *et al*., 2016). Additionally, a wide range of plant species have an extra apoplastic diffusion barrier localized just beneath the epidermis, known as the hypodermis or exodermis (Enstone, Peterson and Ma, 2002). The term exodermis is used when the hypodermis contains a localized lignification and suberin deposition in its cell walls, serving a similar barrier function as the endodermis (Enstone, Peterson and Ma, 2002; Kajala *et al*., 2021; Manzano *et al*., 2022). The exodermis differs from the endodermis in its pattern of differentiation. In maize roots for instance, the CS follows a synchronous development pattern, as ring-like structures, within the entire endodermis. In later stages of development, the CS increase in width, thereby enclosing the entire central cylinder (Enstone, Peterson and Ma, 2002). The SLs are deposited later, but less synchronously, starting from a patchy zone that will develop in a fully suberized endodermis (with exception from the passage cells), depending on the growth conditions (Enstone, Peterson and Ma, 2002; Kreszies *et al*., 2020; Andersen *et al*., 2021; Sexauer *et al*., 2021). In contrast, the development of the exodermis in maize is rather irregular in both radial and longitudinal directions (Líška *et al*., 2016). A recent study has also shown that suberization in the exodermis is essential for survival of tomato under drought conditions, revealing an important physiological function for this cell type (Cantó-Pastor *et al*., 2024). However, we still lack insights on how these two layers are involved in the emergence of the LRP.

Using crop plants for conducting LR studies is a challenging task due to their demanding growth requirements (Garvin, 2007; Scholthof *et al*., 2018). Instead, the wild grass *Brachypodium distachyon* (Brachypodium), possesses several characteristics that make it an excellent monocot model for studying LR development (Raissig and Woods, 2021). Brachypodium has a relatively small genome size, simple growth requirements, fast regeneration time, and exhibits self-pollination. Its embryonic root anatomy consists of a single axial primary root with seminal and leaf node roots developing later depending on the growth conditions. The general radial organization of the primary root consists of an epidermis, five cortex layers, and a single endodermis (Hardtke and Pacheco-Villalobos, 2015). The stele is surrounded by a single pericycle, and the vasculature is arranged with alternating xylem and phloem poles. Most of the above features are closely similar to root anatomy described in major cereal crops (Chochois, Vogel and Watt, 2012; Hardtke and Pacheco-Villalobos, 2015; Raissig and Woods, 2021) including the site of LR initiation (Yu *et al*., 2016). For instance, in rice, maize and barley LRs initiate from cell divisions in pericycle cells associated to the protophloem, so-called phloem pole pericycle cells (Jansen *et al*., 2013; Ni *et al*., 2014; Xiao *et al*., 2019). Thus, the Brachypodium root system exhibits a high degree of developmental and anatomical similarity to important cereal crops, but with less complexity.

In this study, we present a developmental atlas describing the developmental stages of LR development in Brachypodium. We show that the endodermis reactivates its cell cycle and appears to contribute to the formation of the root cap and the formation of columella cells of the emerged LRs. Furthermore, our results indicate the auxin signaling as reported by DR5 promoter activity is not evident in the phloem pole pericycle and during the early stages of LR development. Instead, auxin responses rather appear to be correlated with cell wall modifications during the emergence of the LRP. We show that early suberin deposition in roots appear to be controlled by water and nutrient availability and LRPs emerge towards the growth medium. Finally, we show that the lignification pattern in the exodermis suggests a possible role in the timing of LRP emergence.

## RESULTS

### Lateral roots initiate from phloem pole pericycle cells in Brachypodium

To characterize the sequential developmental stages during LR development in the Brachypodium accession Bd21-3, we adapted the DEEP-CLEAR (Pende *et al*., 2020) protocol for plant tissue to clear roots and used propidium iodide (PI) to visualize the nuclei of the cleared roots via multiphoton microscopy (Figure 1, Figure S1). To categorize the LRP development in Brachypodium, we used the model described for Arabidopsis (Malamy and Benfey, 1997) with a few adaptations in the later developmental stages:

**Figure 1.**
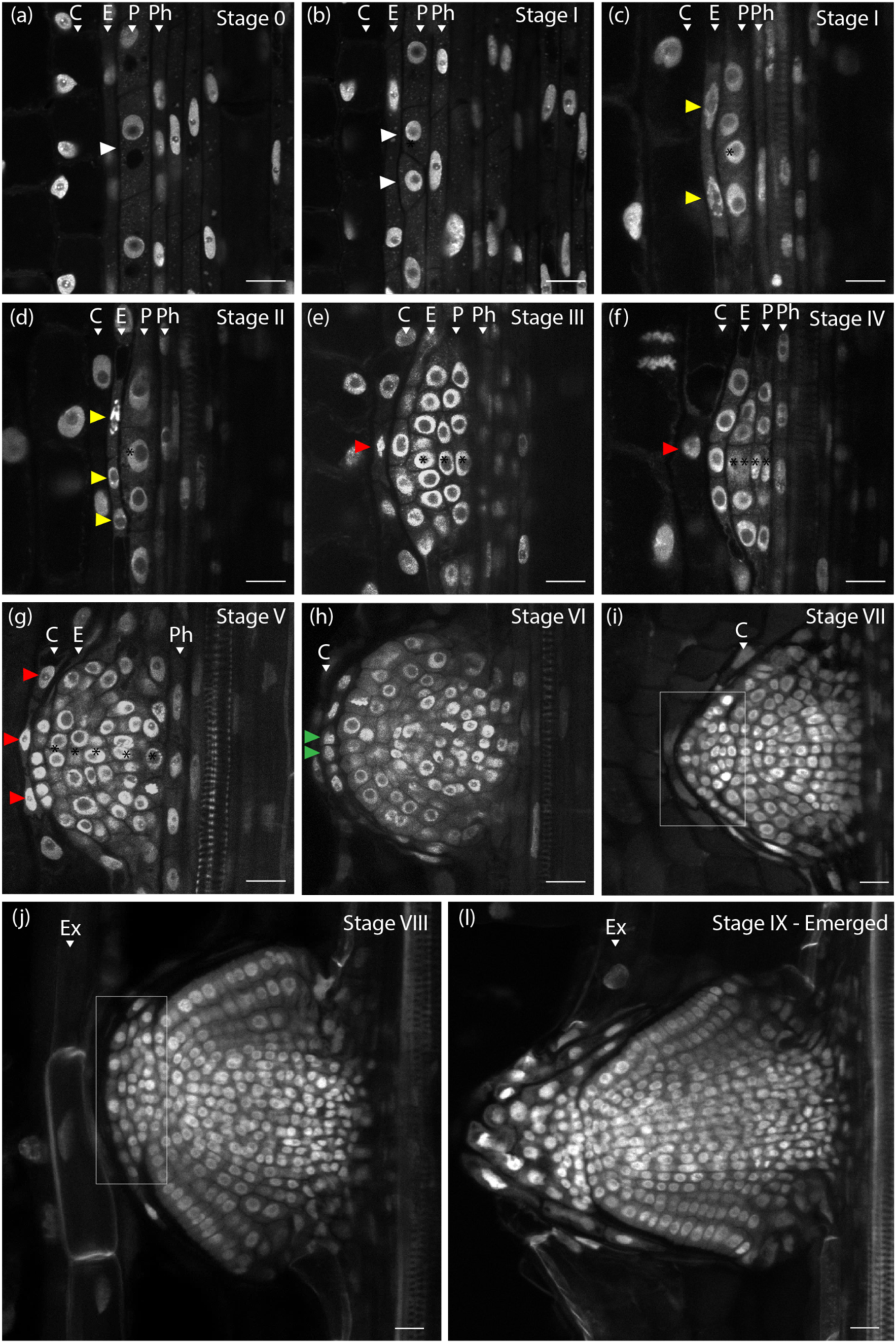
Different stages of LRP development in Brachypodium. (**a**) Stage **0**: No discernible cell divisions in the pericycle cells. (**b**) Stage **I**: White arrows indicate the first anticlinal cell division in the pericycle. (**c**) Stage **I**: Yellow arrows indicate the flattening of the endodermal cells preceding the cell divisions in the next stage. (**d**) Stage II: The endodermis starts to divide anticlinal (yellow arrows). (**e**) Stage **III**: Periclinal divisions take place at the center of the LRP resulting in three layers of cells. The red arrow indicates cell divisions in the overlying cortex. (**f**) Stage **IV**: The LRP undergoes radial expansion through constant anticlinal and periclinal cell divisions in the center of the LRP. Four pericycle cell layers can be observed. (**g**) Stage **V**: Five to six pericycle cell layers can still be distinguished. LRP boundaries are established, and the endodermis appears to become integrated in the LRP. Red arrows indicate more cell divisions in the cortex layer in the vicinity of the LRP. (**h**) Stage **VI**: The endodermal cells on the apex of the LRP start to divide again (green arrows). Pericycle cell layer counting is no longer used from this stage. (**i**) Stage **VII**: Formation of the root cap (white rectangular area. (**j**) Stage **VIII**: The LRP reaches the root exodermis. (**l**) Stage **IX**: Emerged: The LRP is fully formed and traverses the exodermis and epidermis. (**Ex**) Exodermis, (**C**) Cortex, (**E**) Endodermis, (**P**) Pericycle (**Ph**) Phloem. Scale bar = 20 μm.

**Stage I**: Cell divisions occurring in pericycle cells adjacent to the phloem poles (between two xylem poles) are the first anatomical signs of LR initiation (Figure 1b,c, Figure S1b).

**Stage II:** This stage is marked by the initiation of the first anticlinal cell divisions in the endodermal cells overlying a stage I LRP. Subsequently, the cells within the LRP undergo periclinal while endodermal cells continue to divide anticlinal (Figure 1d, Figure S1c).

**Stage III:** Periclinal divisions occur in the center of the LRP resulting in the formation of three layers of cells. In parallel, the endodermis continues to divide anticlinal forming a boundary that span the entire LRP. Additionally, the innermost cortex cell layer, in contact with the endodermis, appears to flatten (Figure 1e, Figure S1d).

**Stage IV:** This stage is characterized by ongoing radial expansion of the LRP through additional anticlinal and periclinal divisions in its central region (Figure 1f). As a result, the canonical dome shape begins to become apparent, and four layers of cells can be counted. Furthermore, ongoing anticlinal cell divisions in the first cortex cell layer were observed, although these cells did not appear to become incorporated in the LRP (Figure 1f).

**Stage V**: Five distinct cortex cell layers remain discernible. The LRP boundaries become well-defined, and the cell divisions patterns indicate that the endodermis-derived cells are integrated within the LRP (Figure 1g, Figure S1e).

**Stage VI:** In this stage the first periclinal endodermal cell divisions occur at the apex of the LRP suggesting the initiation of the lateral root cap formation (Figure 1h, Figure S1e). On average, LRP still have six to seven layers of cells. However, due to the increasing number of cell divisions in the central part of the LRP, it is impossible to apply the cell layer counting system for the remaining LRP developmental stages.

**Stage VII**: The LRP resembles a mature root tip containing an early developmental stage of the lateral root cap that continues to divide anticlinal (Figure 1i, Figure S1f)

**Stage VIII**: At this stage the LRP reaches the exodermis.

**Stage IX**: The LRP crosses the exodermis and epidermis characterizing its full emergence towards the root surface. Figure S2 shows an illustration for the LRP developmental stages in Brachypodium.

### Auxin responses are not detected in the LRP during early stages of development

The phytohormone auxin plays a crucial role during all stages of LRP development in many plant species including important cereal crops such as rice (Lin and Sauter, 2019), maize (Yu, Hochholdinger and Li, 2019) and barley (Kirschner *et al*., 2017). However, most of the insights on how auxin signaling coordinates LR development comes from studies in Arabidopsis (Fukaki and Tasaka, 2009; Vermeer *et al*., 2014; Guseman *et al*., 2015; Cavallari, Artner and Benkova, 2021), and much less is known whether discrete auxin-driven developmental modules have a similar role in monocots.

To characterize transcriptional responses to auxin during LR development, we utilized a *DR5pro::ER-mRFP* marker line (van der Schuren *et al*., 2018). In contrast to what was described for LR initiation in maize (Jansen *et al*., 2013) we were unable to observe a clear DR5pro::ER-mRFP signal in phloem pole pericycle cells and in **Stages I**-**II** LRP (Figure 2a,b); thus, making it challenging to correlate tissue specific changes in auxin responses with LRFC specification and LR initiation. We did observe a weak DR5pro::ER-mRFP signal in the endodermis and most inner cortex cells overlying stage I-II LRP (Figure 2a-b). The earliest detectable DR5pro::ER-mRFP signal in the LRP was only observed at **Stage III**, when the endodermis is already actively dividing (Figure 2c). In later stages, the DR5pro::ER-mRFP signal was no longer detected in the endodermis but it gradually intensified at the apex of the growing LRP (Figure 2d). As the LRP developed (Stages **IV** to **VIII**), we observed an increased DR5pro::ER-mRFP signal in the cortical cell layers overlying the developing LRP (Figure 2d-e). Prior to and after to emergence, the DR5pro::ER-mRFP signal in the newly-formed LR exhibited an expression pattern comparable to the tip of the main root (Figure 2e-g and Figure S4). Notably, we also observed a clear DR5pro::ER-mRFP signal in the exodermis cells overlying the LRP (Figure S4). Although we have observed that Brachypodium LR development is induced under auxin treatment (Figure S6), we failed to detect a DR5pro::ER-mRFP signal in **stage I** LRP. Based on these observations, we characterized whether SISTER of PIN-FORMED 1 (SoPIN1) and AUXIN RESISTANT 1 (AUX1), known transporters involved in auxin efflux and import, respectively, were expressed in early stage LRP (Marchant *et al*., 2002; Reinhardt *et al*., 2003; O’Connor *et al*., 2017). Although we were unable to detect DR5pro::ER-mRFP during LR initiation, the presence of SoPIN1-Citrine was evident as early as **Stage I** (Figure S5a). Later, signal was observed in the endodermis coinciding with its initial cell divisions in the endodermis **Stage II** (Figure S5a). Subsequently, in later stages, SoPIN1-Citrine expression became predominantly localized in the central region of the LRP (Figure S5a). Conversely, AUX1-sGFP exhibited expression within the LRP starting from **Stage I** initially in the phloem-pole pericycle and in the flanking regions with its intensity increasing subsequently in both the vasculature and endodermis (**Stages III** to **V**). Robust expression within the vasculature was also observed in **Stage V** and **VI (Figure S5b)**.

**Figure 2.**
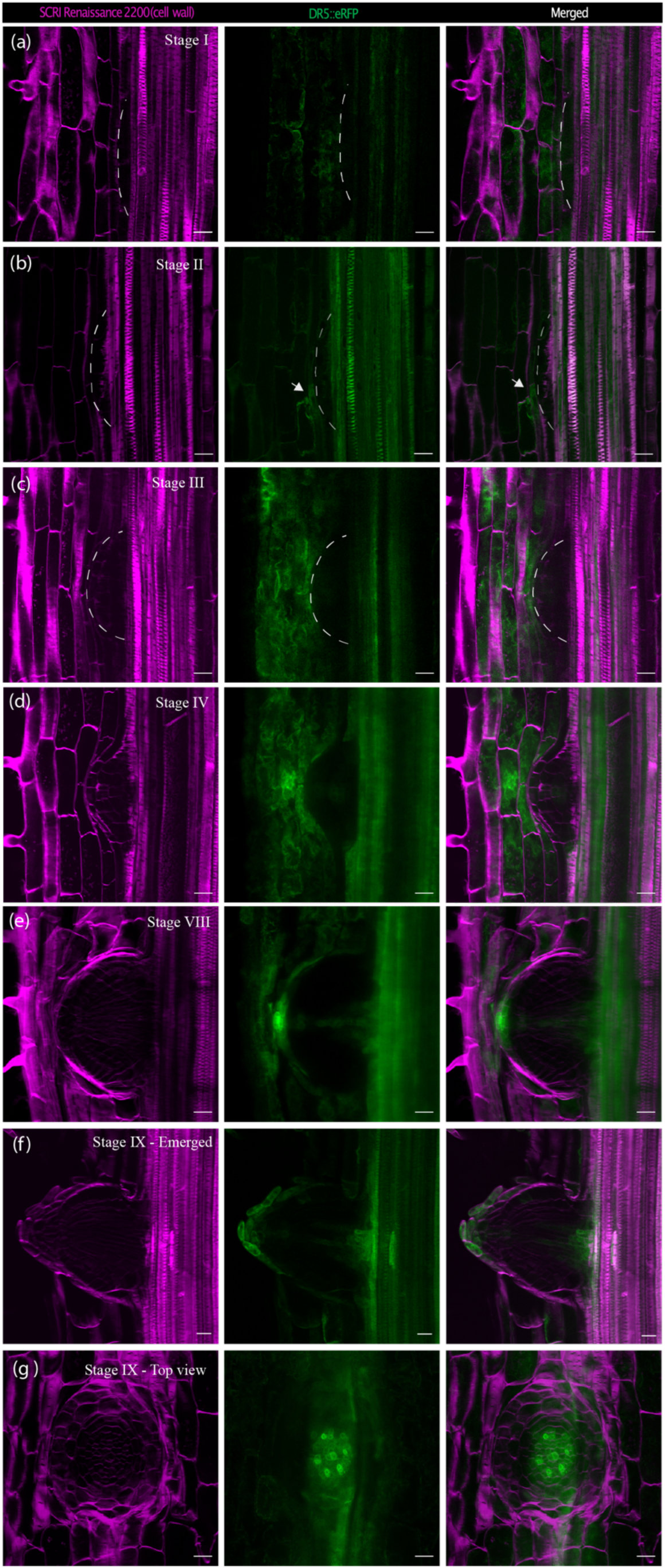
DR5pro::ER-mRFP activity during LR development in Brachypodium. (**a**) The DR5 signal is not evident in Stage I during the first pericycle cell divisions. (**b**) The DR5 signal could be observed in Stage II when the endodermis starts to divide (white arrowheads). (**c**,**d**) The DR5 signal is no longer observed in the endodermis but in the cortex cell layer in the vicinity of the LRP and in the central part of the LRP resembling vasculature. (**e**) The DR5 signal is intensified at the apex of the LRP, in the vasculature, and in the last cortex cell layer. (**f,g**) A fully emerged LR shows a similar DR5 pattern as usually observed in the primary root. Scale = 50 μm.

### Endodermal-derived cells form the root cap of new LRs

Next, we investigated whether the endodermal cells that reactivated their cell cycle and underwent anticlinal and periclinal divisions contribute to the formation of the columella of the LRP. Starch granule formation serves as a marker for differentiation of the columella cells (Guyomarc’h *et al*., 2012; Roychoudhry *et al*., 2023). To assess columella formation, we utilized Lugol’s staining in conjunction with our histological clearing approach. Columella cells (Boxed area in Figure 6) were characterised by the presence of sediments of amyloplasts. Even though the endodermis starts to undergo periclinal divisions from **Stage V** (Figure 6b-c), starch accumulation was only observed in the first cell layer of the columella during late **Stage VI-VII** (Figure 6e-f), following numerous rounds of periclinal cell divisions. The intensity of Lugol’s staining gradually intensified from **Stage VII** to the fully emerged LRP (Figure 6h).

### Water and nutrient availability influence early suberin deposition in Brachypodium roots

During the growth of Brachypodium seedlings on plate, we observed a very strong hydropatterning effect (Figure S3) (Orosa-Puente *et al*., 2018). Basically, all LRs emerged on the side of the root in contact with the growth medium. It is proposed that hydropatterning serves to ensure roots have access with to water and nutrients (Möller, Xuan and Beeckman, 2017). The regulation of the deposition of the SL in the cell wall of endodermal cells has been shown to be responsive to the water and nutrient status (Barberon *et al*., 2016). Moreover, LR formation in Arabidopsis was shown to be accompanied by dynamic changes in suberizaton status of overlying endodermis cells (Ursache *et al*., 2021). In addition, Brachypodium, like many monocots, has an additional cell layer that undergoes localized suberin deposition, the exodermis (Sexauer *et al*., 2021) Using Fluorol Yellow (FY) staining of cleared roots, we confirmed that the pattern of endodermal and exodermal suberization in Brachypodium also occurs after CS establishment initially with patchy zones for both the endodermis and exodermis (Figure 4). This observation is consistent with the findings reported in rice, barley and tomato (Cai *et al*., 2011; Líška *et al*., 2016; Cantó-Pastor *et al*., 2024). Interestingly, suberization in the exodermis appeared delayed compared to suberin deposition in the endodermis (Figure S7). As previously reported in maize, Brachypodium, when grown vertically on agar plates, shows that endodermis and exodermis cells closest to the growth medium are the last to deposit suberin (Figure 4).

**Figure 3.**
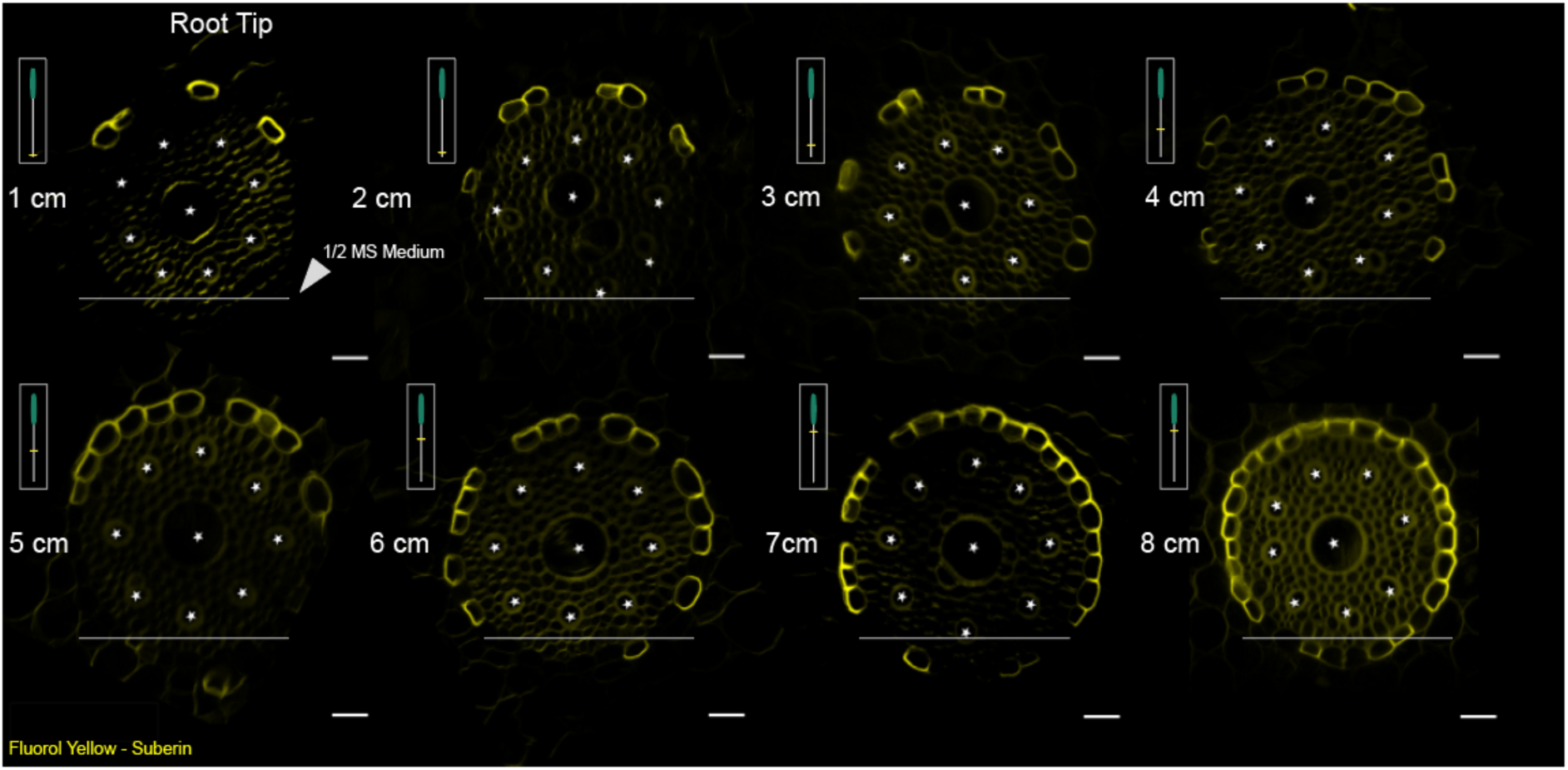
Pattern of endodermal suberization along the Brachypodium root axis adjacent to the nutrient medium growth. Cross-sections of Brachypodium primary root showing asymmetric suberization. SL were developed unilaterally on the side of the root exposed to the air from the root apex, but not on the side exposed to nutrient medium. Scale = 20 μm.

**Figure 4.**
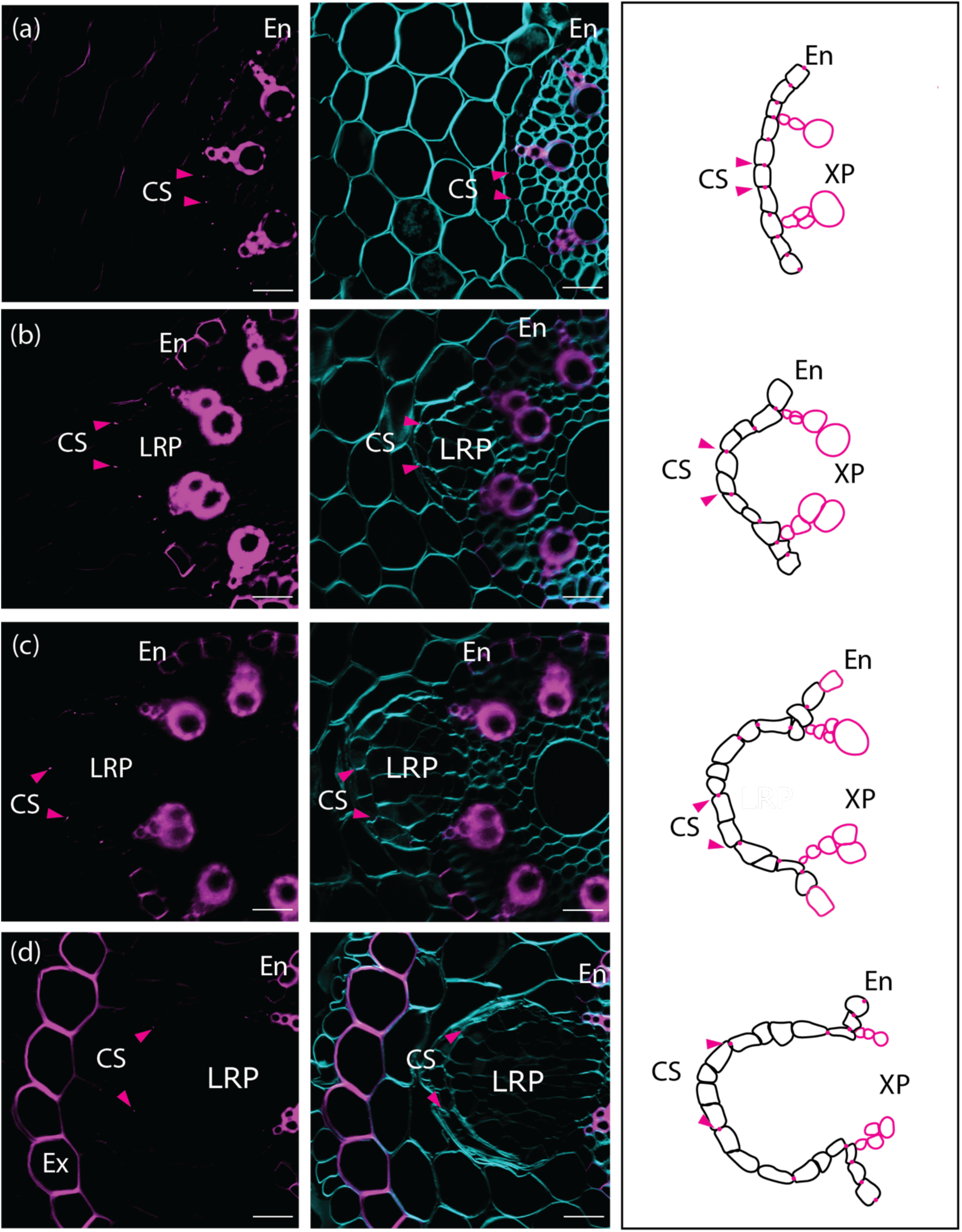
Recently divided endodermal cells do not establish Casparian strips. (**a**-**d**) Cross-sections stained with BF (lignin) in magenta and Renaissance SR2200 (cellulose) in cyan. The images illustrate the progression of cell divisions in the endodermis and the distancing of the previously formed CS. (left) A graphical representation depicting the cell divisions in the endodermis and the separation of the CS is shown on the left. The arrows indicate the position of the CS. Scale = 50 µm.

### Dividing endodermal cells overlying the LRP appear not to establish a Casparian strip

The participation of the endodermis during LR organogenesis in Brachypodium is not unique, as it has been demonstrated already for a range of plant species (Xiao *et al*., 2019). Although we could not observe suberin deposition in the endodermis overlying the LRP under our growth conditions, the Casparian strip domain (CSD) and CS are already present in the overlying endodermis prior to LR initiation. However, little is known regarding the cell fate of the endodermis cells that reactivate their cell cycle and eventually become a part of the LRP. Do these cells after division establish a CSD that is attached to the CS? To address this, we used a histochemical staining for lignin (Basic Fuchsin) and cellulose (Calcofluor White) to counterstain cell walls. In root sections containing LRP, we observed that endodermal cells that underwent anticlinal divisions appear not to establish a CSD, similar to endodermal cells undergoing periclinal divisions, based on the absence of the characteristic lignified spot in the endodermal cross wall (Roppolo *et al*., 2011) (Figure 5). From surface projections of root sections containing LRP in which the endodermis already underwent a few rounds of divisions, it appeared that no newly established CS were present in these endodermis cells (Figure 5b-d). We observed that the CS appeared to undergo a regulated breaking like what was observed during Arabidopsis LR formation (Vermeer *et al*., 2014). In addition, we observed that the CS appears to undergo a lateral detachment (“sliding”) to facilitate the outgrowth of the LRP (Figure S8).

**Figure 5.**
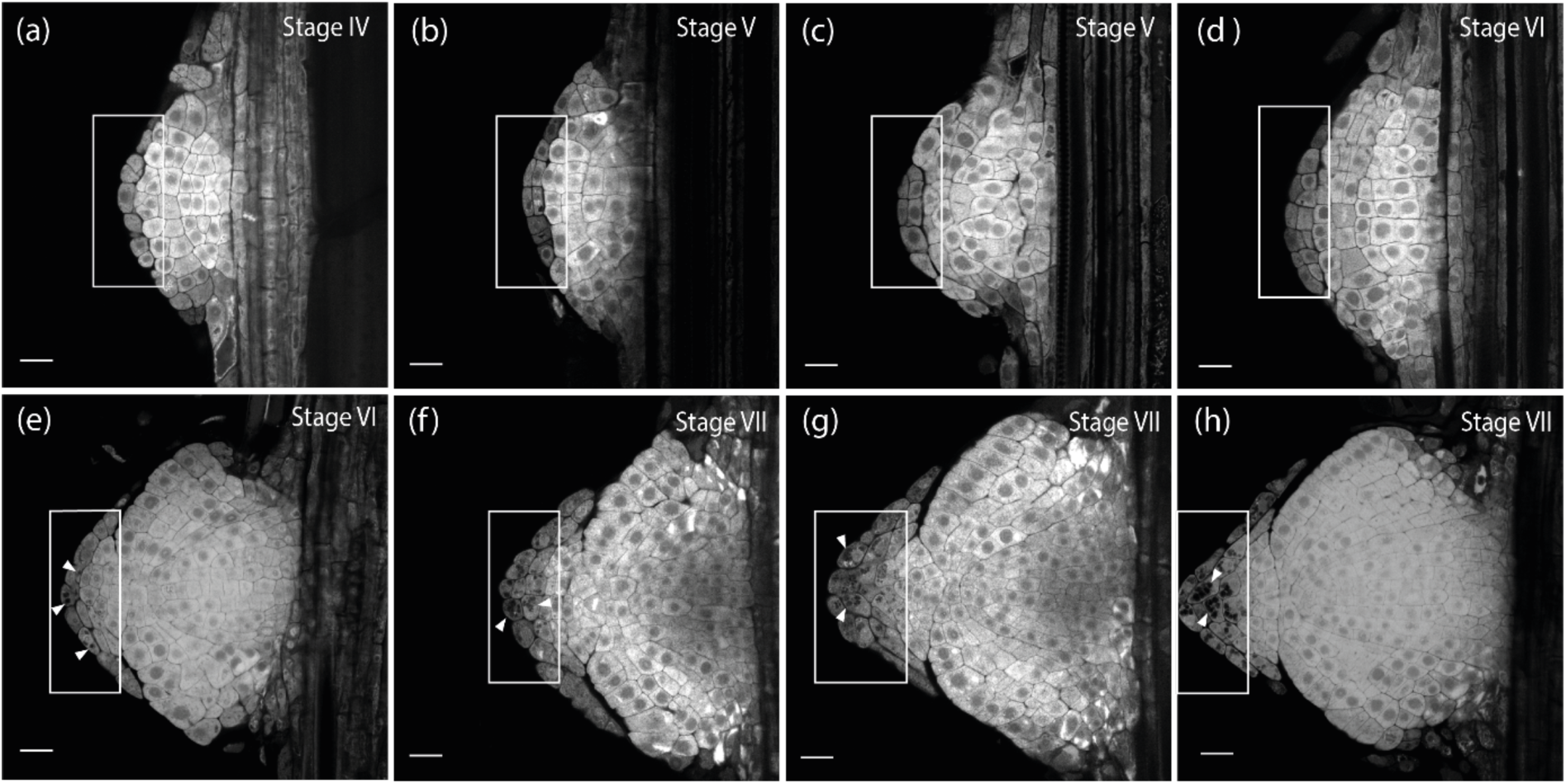
Endodermal cells give rise to the columella cells of the root cap. Starch granules (dark structures in boxed area) were detected using Lugol staining. The boxed area shows cell divisions in the endodermis and its progression in differentiating into columella cells from Stage VI (**e**) to Stage VII (**h**) marked by the sediments of starch granules (arrowheads) in the first cell layer on the apex of the LRP in Stage VI. Scale Bar 50 µm.

## Discussion

In this study, we present an atlas describing the consecutive stages of LR development in Brachypodium based on the model utilized for Arabidopsis (Malamy and Benfey, 1997; Péret, Larrieu and Bennett, 2009; Van Norman *et al*., 2013; Vermeer and Geldner, 2015; Wachsman and Benfey, 2020). We show that in Brachypodium, like other monocots, pericycle cells adjacent to phloem are competent for LR organogenesis. Furthermore, the endodermis undergoes mitotic activation soon after the initial pericycle cell divisions and overlying endodermal cells will become an integral part of the LRP in later developmental stages, as they will form the root cap (Figure 1 and 6). While the inner-most cortex layer appears to undergo cell divisions, we could not confirm whether these cells become part of the LRP itself, like the overlying endodermal cells. Alternatively, we hypothesize these divisions are required to accommodate the expansion growth of the LRP, thereby facilitating emergence. Although most of textbook knowledge regarding LR development is based on Arabidopsis studies, the mitotic reactivation and participation of the endodermis and derived cells are observed in a large number of plants species including barrel clover (Herrbach, Maillet and Bensmihen, 2018), maize (Jansen *et al*., 2013) barley (Orman-Ligeza *et al*., 2013) and many others (Xiao *et al*., 2019). It appears that absence of the incorporation of the endodermis during LRP growth could be specific for the Brassicaceae (Xiao *et al*., 2019). Alternatively, we hypothesize that these divisions are necessary to accommodate the expansion growth of the LRP, potentially aiding in its emergence by facilitating growth through the overlying both the endodermis and adjacent cortex cell layers (Bell and McCully, 1970; Torres-Martínez *et al*., 2020). The mitotic reactivation of the endodermis raises important questions: What triggers this process and do the dividing endodermal cell change their identity and if so at what stage? At which stage do they obtain columella identity? To address these questions follow-up studies employing sectorial (mosaic) analyses and high-resolution expression analysis (single cell/ nuclei sequencing, spatial transcriptomics) (Birnbaum, 2018; Torres-Martínez *et al*., 2020; Liu *et al*., 2023) would be a logical step to track cell lineages and changes of cell identity during the LRP developmental process.

Auxin serves as a crucial regulator of LR patterning, development and the DR5 reporter is commonly employed to visualize auxin responses dr5 (Ulmasov *et al*., 1997; Liao *et al*., 2015). In this study, we could not detect the DR5pro::ER-mRFP (DR5) signal in phloem pole pericycle cells during the formative cell divisions leading to a stage I LRP. However, there was induction of DR5 signal in the overlying endodermis and even more so in the cortex cells overlying the LRP. Interestingly, we also observed during later stages of LRP development, clear induction of the DR5 signal in the exodermis overlying the LRP. This suggests for a similar role of auxin signaling to regulate cell wall modifications to facilitate the emergence of the LRP (Swarup *et al*., 2008; Meng *et al*., 2019). The observed induction of the DR5 reporter in the overlying exodermis suggest for a possible role for auxin signaling to regulate cellular responses, such as modification of the lignin barrier, to accommodate emergence (Nakayama *et al*., 2017).

Previous studies have reported the absence or presence of a weak DR5 signal during the first cell divisions in the pericycle of rice (Ni *et al*., 2014) barley (Kirschner *et al*., 2017) and maize (Jansen *et al*., 2012)opposite to what is commonly observed in Arabidopsis (Dubrovsky *et al*., 2000; Vanneste *et al*., 2005; Marhavý *et al*., 2013). Similarly, during many stages of Brachypodium embryo development, the DR5 signal was not or barely detected(Hao *et al*., 2021) counterintuitive compared to the observations during Arabidopsis embryogenesis (Mo and Weijers, 2009). The synthetic DR5 promoter contains direct repeats of a medium-affinity biding site for the AUXIN RESPONSE FACTORs (ARF) transcriptional regulators (Ulmasov *et al*., 1997; Boer *et al*., 2014). Therefore, it is likely that only part of the transcriptional response to auxin is reported. The use of higher affinity binding sites could provide a solution to better address the role of auxin during early developmental processes in Brachypodium (Liao *et al*., 2015; Hao *et al*., 2021). The same DR5 reporter has been used to monitor changes in auxin signaling during Brachypodium spikelet formation in the shoot (O’Connor *et al*., 2017). In addition, we observed a clear DR5 signal during later stages of LRP development, including in the cortex and exodermis. A plausible explanation would be that a set of ARFs with reduced affinity for the DR5 promoter are regulating auxin responses during early stages of LR development in Brachypodium. It is clear that auxin can induce LR formation in Brachypodium (Pacheco-Villalobos *et al*., 2013) and important regulators of auxin import and efflux are already expressed in stage I LRP (Figure S5).

The root endodermis serves as apoplastic barrier for the radial transport of water and nutrients to the plant’s vascular system (Barberon *et al*., 2016). To fulfil this role, the endodermis relies on the formation of the lignified Casparian strips. Subsequently, suberin lamellae are deposited as a secondary cell wall modification surrounding the plasma membrane (Barbosa, Rojas-Murcia and Geldner, 2019). Moreover, many plant species have an extra barrier called the exodermis, which also exhibits lignin and suberin deposition (Kajala *et al*., 2021; Liu and Kreszies, 2023). Recent studies have reported that the exodermis functions in the tolerance to abiotic stresses (Cai *et al*., 2011; Kajala *et al*., 2021; Manzano *et al*., 2022; Cantó-Pastor *et al*., 2024). Here, we demonstrate that the daughter cells of divided endodermal cells do not appear to form a CSD in their cross walls (Figure 5). Similar to what was described for Arabidopsis, the CS appears to be detached longitudinally and local breaks appear, likely facilitating the emergence of the LRP. While there is increasing interest in studying the function and formation of the CS in monocots such as rice and maize (Karahara *et al*., 2004; Wang *et al*., 2022), little is known whether the CASPARIAN STRIP DOMAIN PROTEINs (CASP) are degraded during LR development as described for Arabidopsis (Vermeer *et al*., 2014). It will be interesting to test whether CASP/CSD degradation could be a general mechanism that allows for loosening and/or local breaking of the CS in plants during LR emergence. Moreover, Brachypodium has a lignified exodermis (Sexauer et al., 2021).However, under our experimental conditions, the exodermis in many cases showed still little lignification at the time of LR emergence.

## CONCLUSION

Here we provide an atlas describing the various developmental stages of LRP development in Brachypodium and shed light on the potential roles of different cell types and molecular mechanisms involved to facilitate their development. This now provides a perfect starting point to dissect the trajectories of cell types and if there are regulatory mechanisms that could be part of conserved modules for root branching in general. Brachypodium LRP formation provides a beautiful (non-domesticated) plant model to investigate how the endodermis and cortex re-activate their cell cycle and contribute to organogenesis and emergence. In addition, it allows the investigation which (hormonal) signaling pathways are re-wired and which are conserved during developmental processes.

## MATERIALS AND METHODS

### Plant Materials and Growth Conditions

*Brachypodium distachyon* seedlings (Bd-21-3) (Vogel and Hill, 2008) were grown vertically on 0.8% agar supplemented with half strength Murashige-Skoog (MS) pH 5.8 at 22° C under long day or constant light. 5 DAG seedlings were collected for analysis. After removal of the seed husk, seeds were surface sterilized using sodium hypochlorite 5% and 0.01% Triton for 4 min and rinsed at least four times in autoclaved desilted water. Seeds were placed on medium (prepared as described above) with embryo towards the bottom of the and facing the lid of the 120mm square plate in order to prevent shoots and roots growing into the media or in the wrong direction. Plates were placed into growth conditions at an angle of about 20° to ensure that roots grow on the medium and not into the air as described by (van der Schuren *et al*., 2018). After a maximum of 7 days in 22 C under long day or constant light, seedlings were harvested for clearing and analysis.

### Auxin Treatment

Seedlings of Brachypodium Bd-21-3 were grown vertically under long day for 5 days on standard ½ MS plates 0.8% agar and then transferred to 10 µM IAA-treated plates. Images were taken at 0, 3, and 7 days after the seedlings were transferred to auxin.

### *Chemicals* for clearing and staining solutions

The following chemicals were used in the DEEP-Clear (Pende *et al*., 2020) adapted version to plant tissues: PFA (paraformaldehyde) (CAS no. 30525-89-4, Merck, http://www.merck.com/), xylitol (CAS no. 87-99-0, Sigma, http://www.sigmaaldrich.com/), urea (CAS no. 57-13-6, Sigma), SR2200 (Renaissance Chemicals), Basic Fuchsin (CAS no. 58969-01-0, Sigma), THEED (Sigma-Aldrich, 87600-100ML), 5% (v/v) Triton X-100 (Roth, 3051.2).

### Preparation of hand-sectioned root samples

For sectioning, roots fragments of 1 cm with the region of interest were embedded in 4% agarose. After solidified, agarose blocks containing the region of interest were glued on a hand microtome and sections of approximately 50 μm were prepared for clearing or immediate visualization.

### Clearing and staining

Clearing steps using DEEP-Clear were performed as described for ClearSee in (Kurihara *et al*., 2015) and adapted from (van der Schuren et al., 2018) for *Brachypodium* samples. DEEP-clear solution consists in 5 to 8% (v/v) THEED, 5% (v/v) Triton X-100, and 25% (w/v) urea in water. Heating the solution is not recommended. 7 DAG old root seedlings were collected for clearing for full root treatment and for semi-thin sectioning. Samples were fixed for 1 hour in 4% (w/v) paraformaldehyde in 1× phosphate-buffered saline (PBS) with 3 rounds of soft vacuum infiltration. After, roots were washed five times in 1x PBS with another round vacuum to ensure the removal of PFA. Samples were then transferred to DEEP-Clear solution for clearing. Fixed root tissue was incubated at room temperature with gentle shaking and solution for 7-10 days and solution replaced twice. For staining of the fixed and cleared tissue, 1% stock solution of Basic Fuchsin (for lignin staining), Fluorol Yellow (for suberin staining), Renaissance and/or Calcofluor (for cell wall staining) were separately prepared directly in DEEP-Clear and stored at 4°C. Working solutions were prepared as in (Ursache *et al*., 2018). In order to combine multiple dyes, samples were incubated first in Basic F (0,1% in DEEP-Clear) for one 1 hour and washed in DEEP-Clear overnight. After, several rounds of washings, samples were transferred to Renaissance (0,1% in DEEP-Clear) for 2 days and washed overnight in DEEP-Clear. Finally, samples were transferred to Fluorol Yellow (FY) (0,01% in DEEP-Clear) for 1 hour and counterstained in aniline blue (0,5% in water) for 1 hour in the darkness. Samples prior FY staining can be stored in 50% glycerol at 4%. FY solutions and FY-stained samples were kept in darkness to prevent bleaching.

### Starch staining

To observe starch granules in the LRPs of Brachypodium, the root cortex was mechanically removed without damaging the LRPs. Roots were then cleared for 3 days in DEEP-Clear. After, roots were dipped in Lugol’s staining solution (Sigma-Aldrich) for 5 minutes, washed with distilled water, and observed under 2-photon microscopy.

### Microscopy

Roots were observed using Leica TCS SP8-MP equipped with a resonant scanner (8 kHz) using 25x, 40x and 63x water immersion objectives. Figures were arranged in Adobe Illustrator (Adobe Systems Inc., http://www.adobe.com/) or in PowerPoint (Microsoft Corporation) and the brightness was increased equally, without further modifications. The 3D reconstruction was done using the Fiji package (Schindelin *et al*., 2012).

---

**Supplemental Figure S1.**
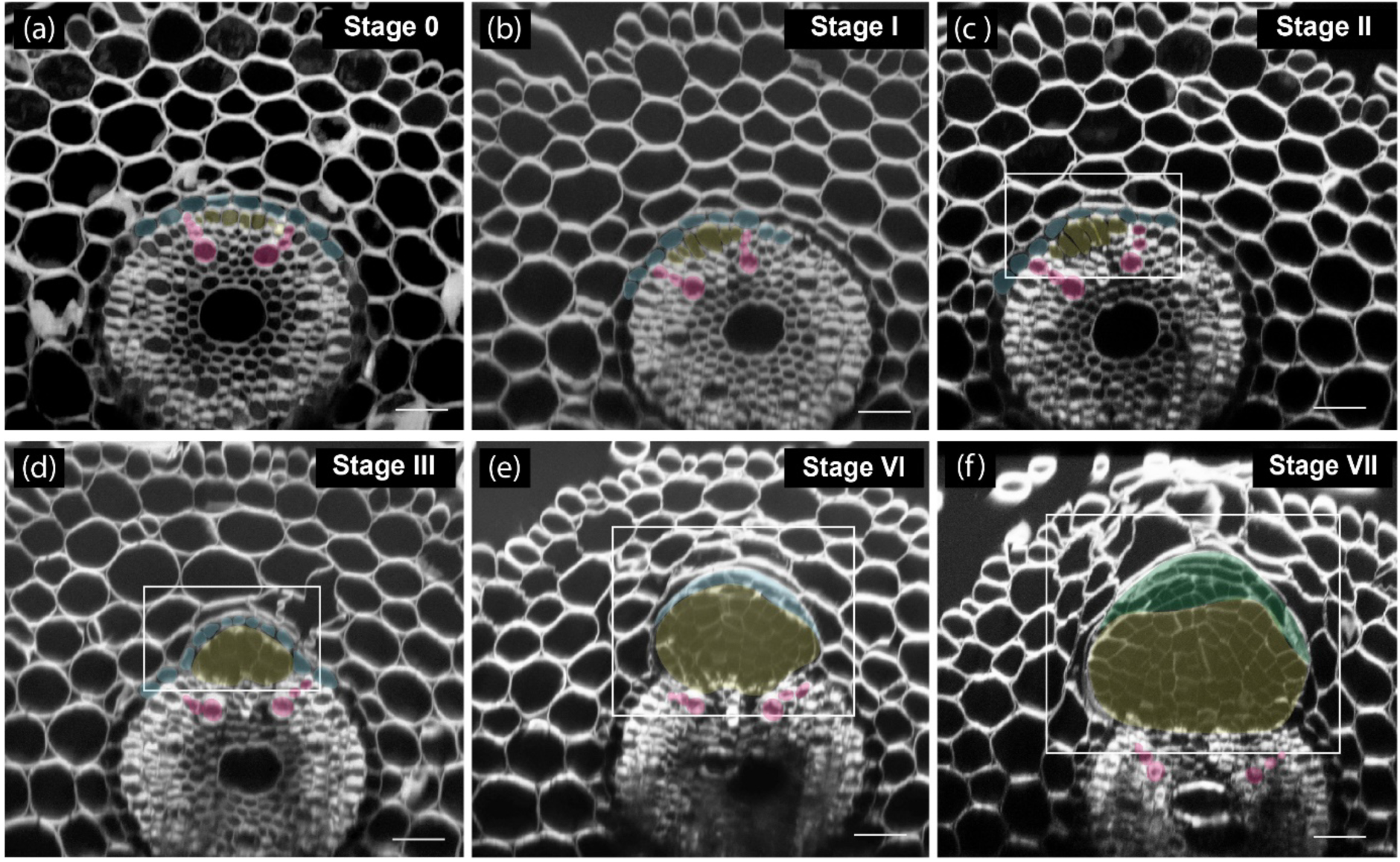
Orthogonal view of developmental stages of LRP formation in Brachypodium. (**a**) Stage **0**: No evident swelling of the pericycle cells (white arrows). (**b**) Stage **II**: Recently divided pericycle cells promote the displacement of the endodermis. In (**c** and **d**) Stage **II** and **III** the pericycle divides anticlinal and periclinal (rectangular area). (**e** and **f**) - Periclinal and anticlinal cell divisions are observed at the apex of the LRP (Stage **V**) following the establishment of the lateral root cap (Stage **VII**). Purple: Xylem, Blue: Endodermis, Yellow: Pericycle, Green: Root Cap. Scale Bar: 50 µm

**Supplemental Figure S2.**
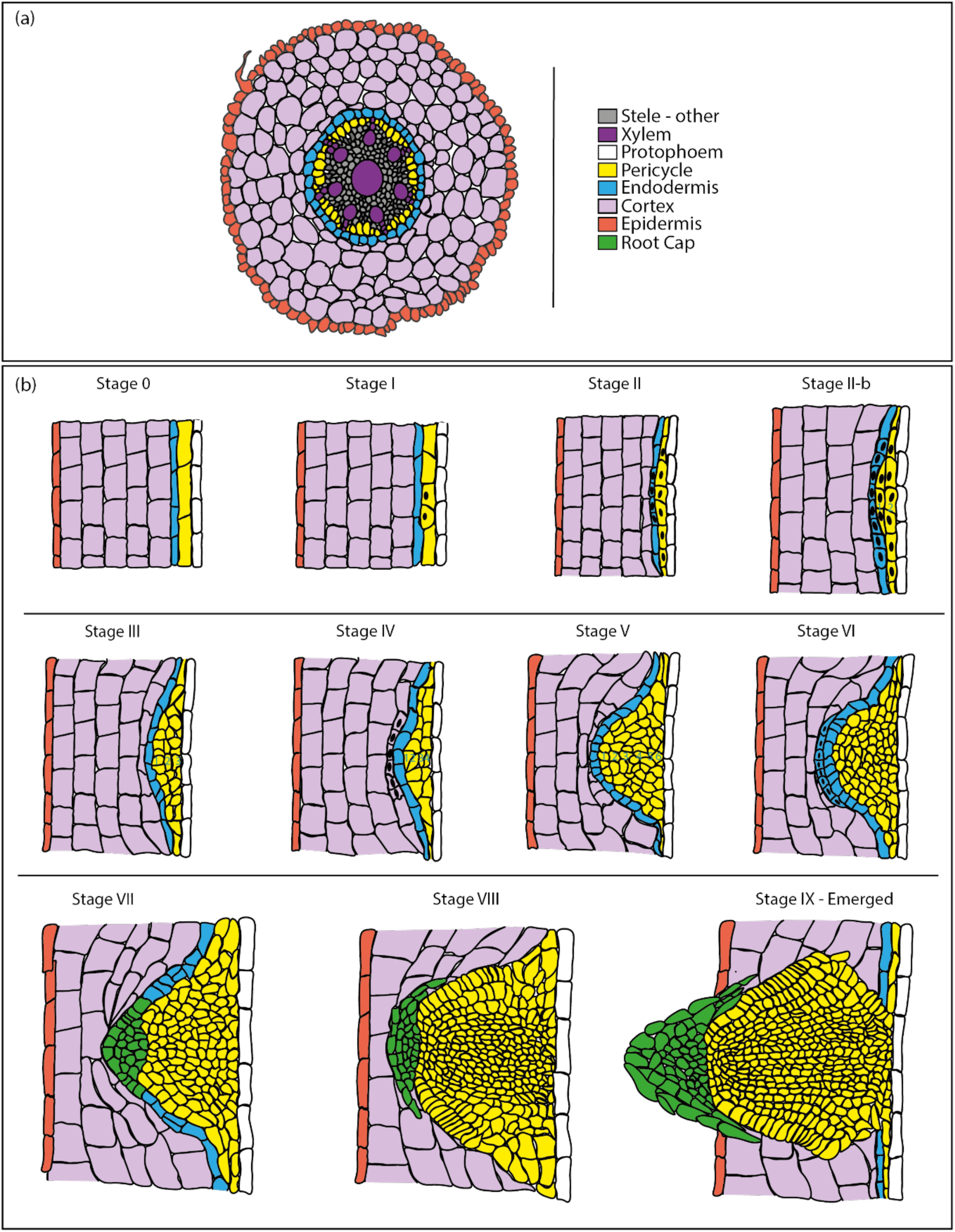
Schematic representation of LRP development in Brachypodium. (**a**) Representation of root cross section of Brachypodium. (**b**) Successive stages of LRP formation are illustrated.

**Supplemental Figure S3.**
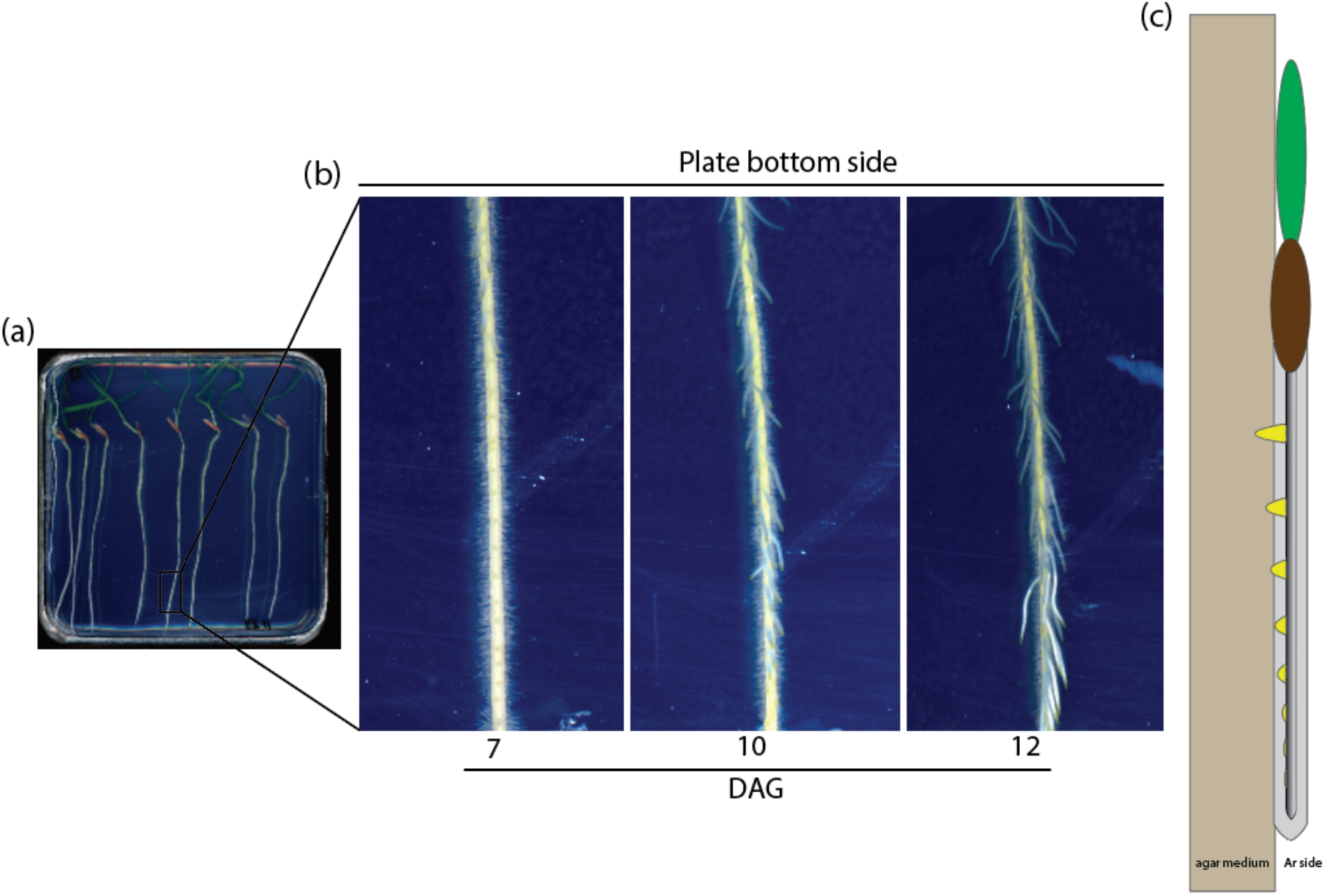
Brachypodium LRs emerge towards the agar medium. (**a**) Brachypodium seedlings grown on 12×12 cm plates supplemented with half-strength MS solution. (**b**) Progression of lateral root emergence towards the agar after 7, 10, and 12 days after germination (DAG). (**c**) Side view illustration of lateral roots growing towards the agar medium.

**Supplemental Figure S4.**
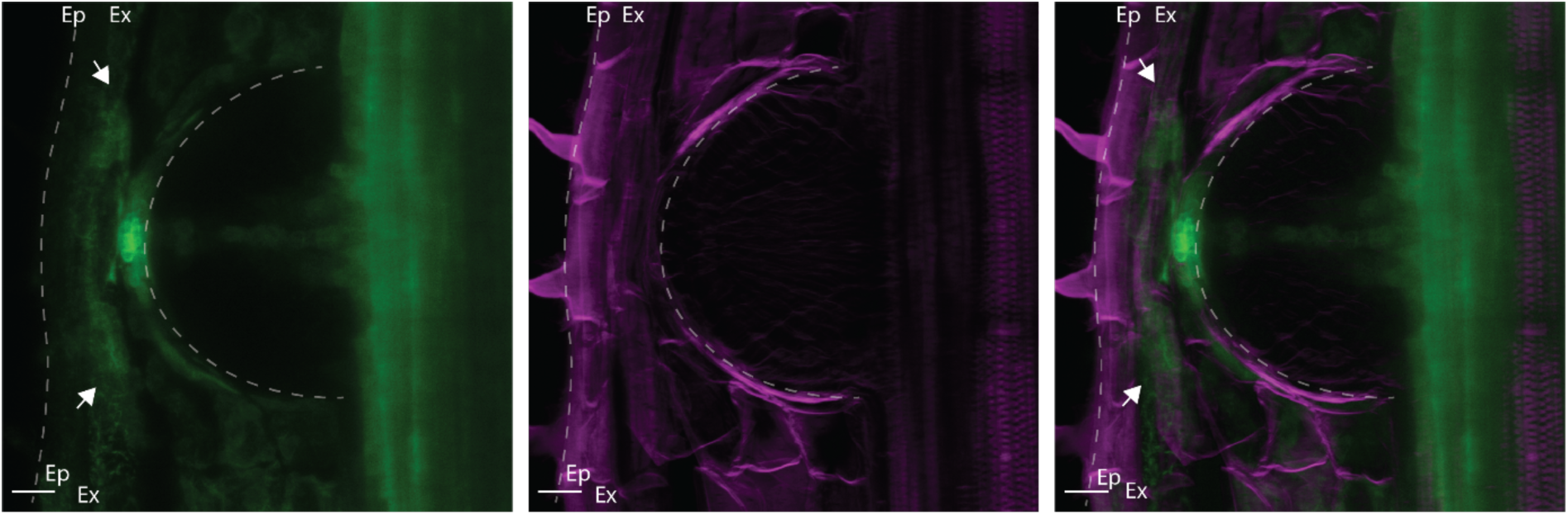
*DR5pro::ER-mRFP* is induced in the apex of the LRP and in the overlying exodermis. The white arrow indicates presence of the DR5 signal (green) confined to the overlying exodermis. Ep: Epidermis, Ex: Exodermis. Scale Bar = 20 µm.

**Supplemental Figure S5.**
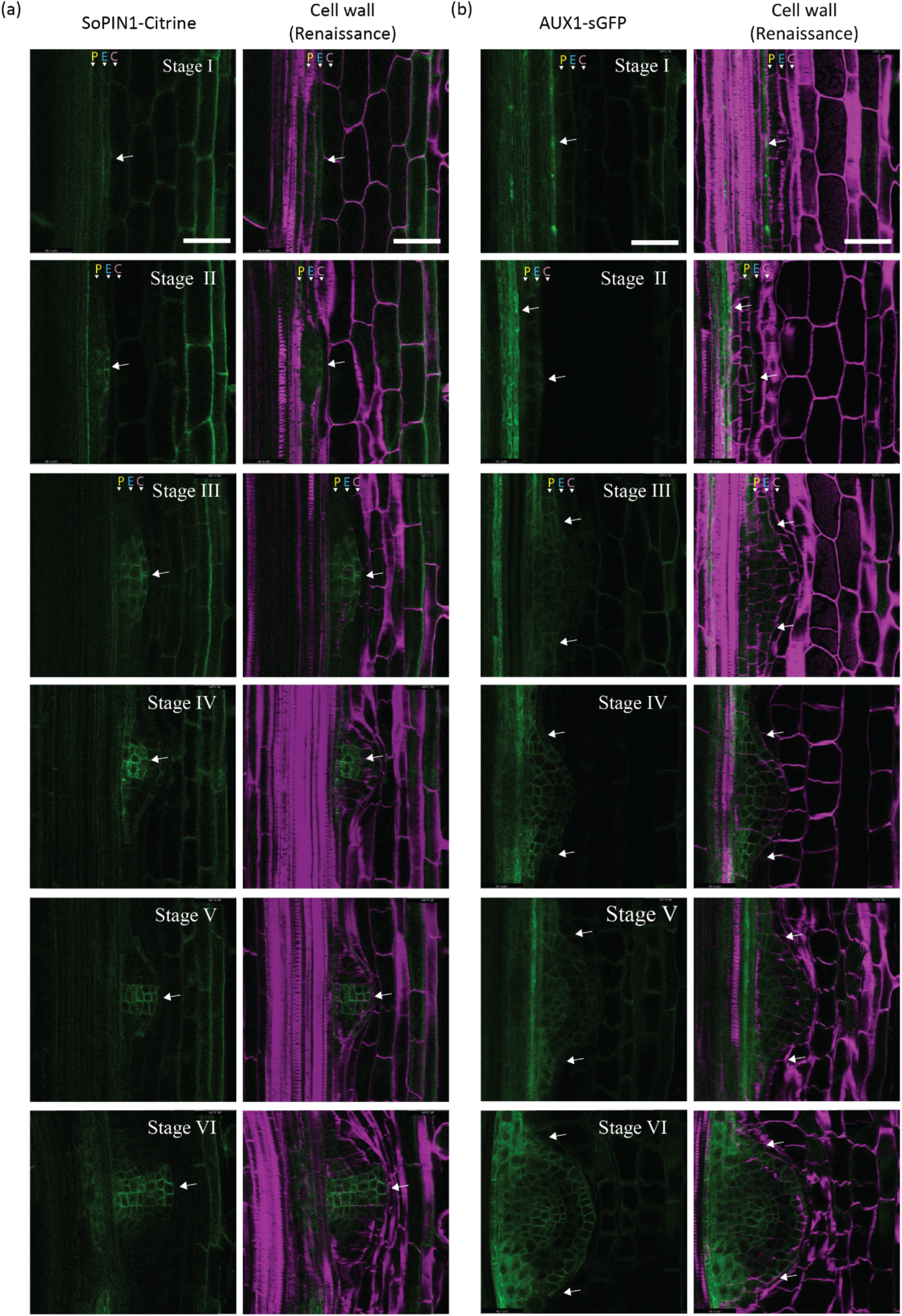
Localization of SoPIN1-Citrine and BdAUX1-sGFP during LR formation in Brachypodium. (**a**) SoPIN1-Citrine is observed from Stage I (white arrowhead) during the initial cell divisions in the endodermis. Later, expression is localized in the central region of the LRP. (**b**) Expression of AUX1-sGFP is observed in the vasculature of the primary root and in the LRP from Stage I (white arrowheads). During later stages of LR development, AUX1-sGFP signal is observed in the flanking regions of the LRP with its intensity increasing subsequently in both the vasculature and endodermis in Stage VI. Scale = 50 μm.

**Supplemental Figure S6.**
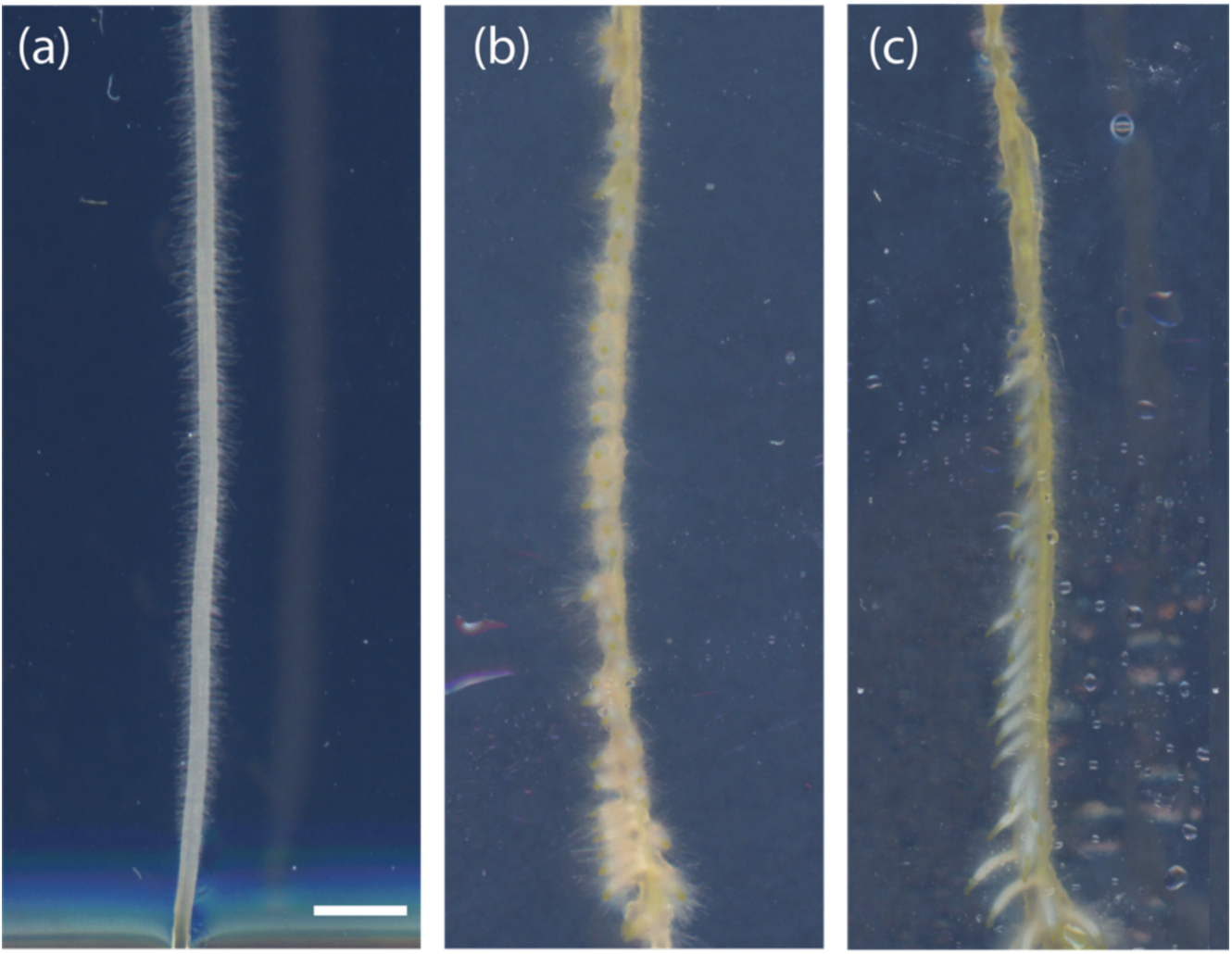
Exogenous auxin addition induces lateral root formation in Brachypodium. Seedlings were grown on standard ½ MS plates and transferred to auxin treatment (10 µM IAA) for 0 (**a**), 3 (**b**), and 7 days (**c**). Scale bar: 0.3 cm.

**Supplemental Figure S7.**
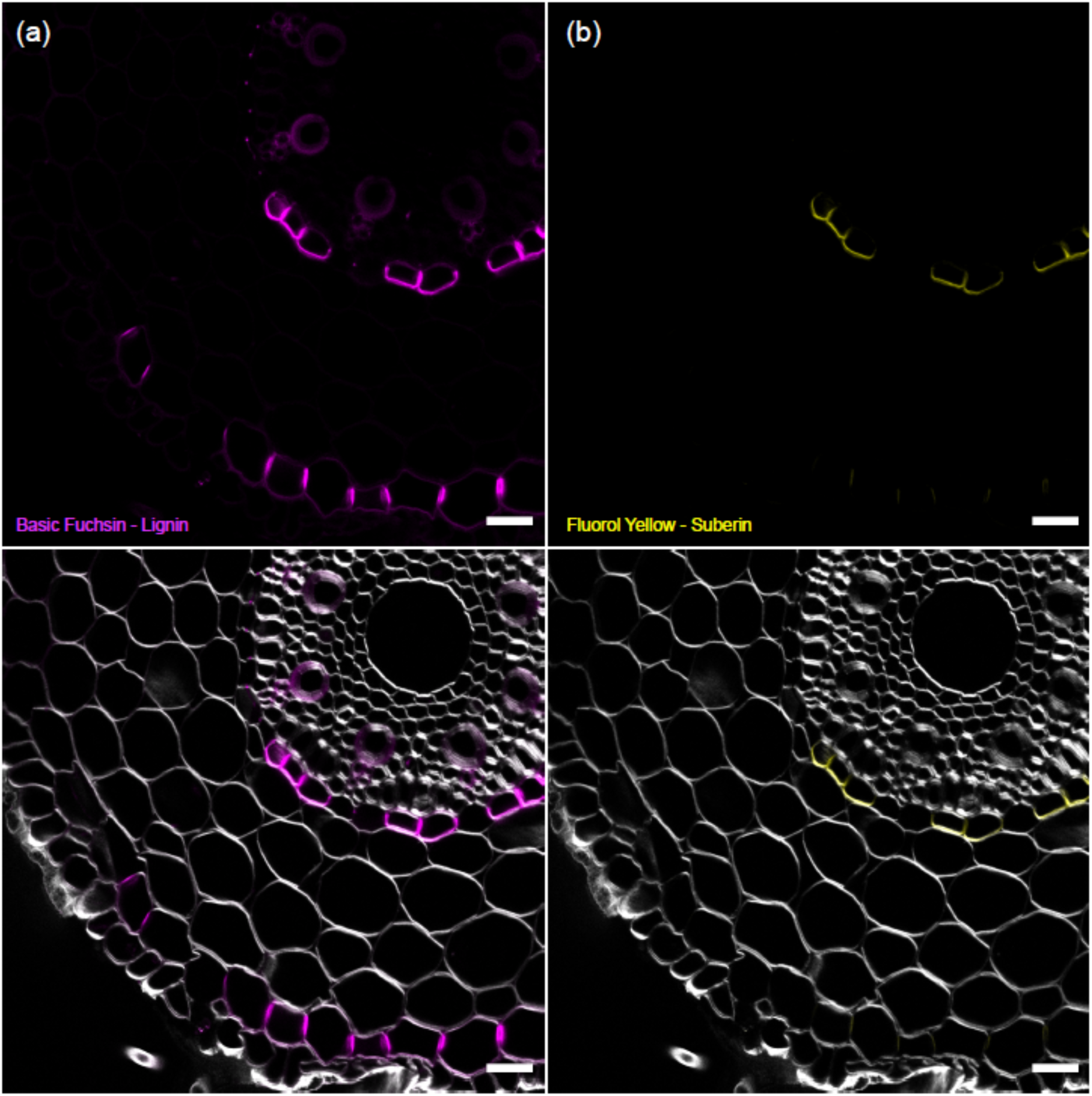
The exodermis show later suberization in comparison to the endodermis. Cross-sections of Brachypodium primary roots double stained for (**a**) lignin (BF) (**b**) suberin (FY) counter stained with Renaissance SR2200 (cellulose). Scale: 50 µm

**Supplemental Figure S8.**
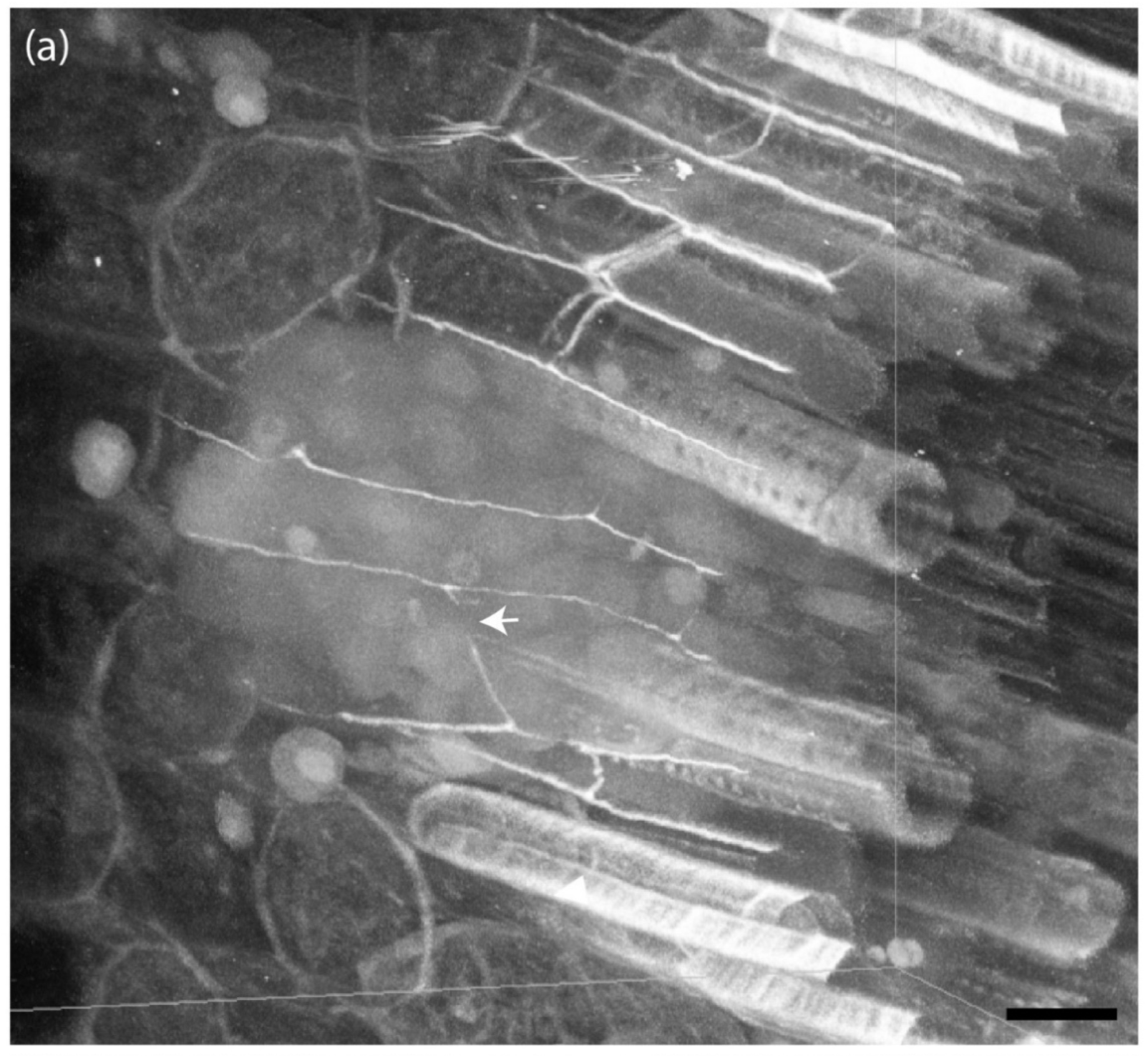
The CS undergoes lateral detachment during LRP development in Brachypodium. Maximum projections of sections stained with BF. The CS is laterally displaced during the emergence of the LRP. Scale bars = 20 µm.

## Notes

### Competing Interest Statement

The authors have declared no competing interest.

